# The porcine deltacoronavirus replication organelle comprises double membrane vesicles and zippered endoplasmic reticulum with double membrane spherules

**DOI:** 10.1101/719443

**Authors:** Nicole Doyle, Philippa C. Hawes, Jennifer Simpson, Lorin H. Adams, Helena J. Maier

## Abstract

Porcine deltacoronavirus (PDCoV) was first identified in Hong Kong in 2012 from samples taken from pigs in 2009. PDCoV was subsequently identified in the USA in 2014 in pigs with a history of severe diarrhea and the virus has now been detected in pigs in several countries around the world. Following the development of tissue culture adapted strains of PDCoV, it is now possible to begin to address questions regarding virus-host cell interactions for this genera of coronavirus. Here we present a detailed study of PDCoV induced replication organelles. All positive strand RNA viruses induce the rearrangement of cellular membranes during virus replication to support viral RNA synthesis, forming the replication organelle. Replication organelles for the *Alpha*-, *Beta*- and *Gammacoronavirus* genera have been characterized. However the structures induced by deltacoronaviruses, in particular the presence of convoluted membranes or double membrane spherules, are unknown. Initially, the dynamics of PDCoV strain OH-FD22 replication were assessed with the onset of viral RNA synthesis, protein synthesis and progeny particle release determined. Subsequently, virus induced membrane rearrangements were identified in infected cells by electron microscopy. As has been observed for all other coronaviruses studied to date, PDCoV replication was found to induce the formation of double membrane vesicles. Significantly however, PDCoV replication was also found to induce the formation of regions of zippered endoplasmic reticulum and small associated tethered vesicles, double membrane spherules. These structures strongly resemble the replication organelle induced by avian *Gammacoronavirus* infectious bronchitis virus.

## 1. Introduction

All positive strand RNA (+RNA) viruses studied to date rearrange the membranes of their host cell to create the virus replication organelle (RO). For several viruses, ROs have been proven to be the site of viral RNA synthesis [1–4]. As such, ROs are thought to facilitate the coordination of the processes involved in viral RNA synthesis, as well as provide an enclosed environment to protect viral RNA from detection by the host cell. This helps to prevent degradation of viral RNA by host enzymes and, importantly, prevent activation of cellular intrinsic antiviral signaling pathways. Therefore RO formation and function are critical steps in the replication cycle of all +RNA viruses. The membrane origin, shape and structure of ROs varies between different virus families but there are common themes in the types of structure produced. Several viruses including enteroviruses, arteriviruses, toroviruses, hepatitis C virus and foot and mouth disease virus induce the formation of double membrane vesicles (DMVs), often along with single membrane vesicles and tubules or paired membranes [5–10]. The second common structure induced are spherules or invaginated vesicles, which are formed by alphaviruses, flaviviruses, bromoviruses and nodaviruses [3, 4, 11–15]. These are small single membrane vesicles pinched out from a cellular membrane but they remain tethered to the membrane and possess a channel connecting the interior of the spherule with the cytoplasm.

An important family of +RNA viruses are the coronaviruses. Coronaviruses infect a wide range of species including pigs, cattle, chickens, cats and humans causing both economically damaging diseases impacting livestock farming industries and important human illnesses. In the last 17 years, two novel coronaviruses, SARS- and MERS-coronaviruses (SARS- and MERS-CoV), have emerged into the human population from zoonotic sources and have caused significant illness with high mortality rates [16, 17]. This, along with the recent emergence of SADS-CoV in pigs arising after a species jump [18, 19], highlights the importance of this family of viruses as a threat to human health with the potential for further emergence of new pathogenic viruses. There are four genera within the coronavirus family, *Alpha*-, *Beta*-, *Gamma*- and *Deltacoronaviruses* and the ROs of viruses within these genera have been characterized. All coronaviruses induce the formation of DMVs. In addition, there are rearrangements of the endoplasmic reticulum (ER) to form either a branching network of membranes referred to as convoluted membranes (CM) found in *Alpha*- and *Betacoronavirus* infected cells [20–23] or paired ER membranes referred to as zippered ER in *Gammacoronavirus* infectious bronchitis virus (IBV) infected cells identified in our previous work [24, 25]. Associated with the zippered ER in IBV infected cells are small double membrane spherules, not seen previously in cells infected with other coronaviruses.

*Deltacoronaviruses* were first characterised as a new coronavirus genus in 2011. The majority of members of this genus infect avian species and have been identified only through sequencing the viral genome. Therefore without viral isolates to able to replicate in cell culture, studying the virus-host interactions of this genus of coronaviruses has not been possible. However, porcine deltacoronavirus (PDCoV) was identified in Hong Kong in 2012 [26] and subsequently from pigs in the USA and other countries [27–31]. The virus causes an acute gastrointestinal infection with severe diarrhea, vomiting and atrophic enteritis [32]. Importantly, cell culture adapted strains of PDCoV have now been developed [33–35] allowing the characterization of how this genus of coronaviruses interacts with its host cell, including a characterization of the *Deltacoronavirus* RO. In a recent publication, Qin *et. al.* confirmed the presence of DMVs in PDCoV infected cells [36]. However, CM or zippered ER and spherules were not identified and the presence of these structures remains to be determined. Here we have characterized PDCoV stain OH-FD22 [34] replication in porcine LLC-PK1 cells, including a detailed characterization of ROs.

## 2. Materials and Methods

### 2.1. Cells and virus

Porcine LLC-PK1 cells (ATCC CL-101) [37] were maintained in DMEM supplemented with 10% FCS. Porcine deltacoronavirus OH-FD22 was kindly provided by Prof. Linda Saif, The Ohio State University [32, 34]. Viral infection of LLC-PK1 cells was performed in EMEM supplemented with 1% HEPES, 1% NEAA and 1% antibiotic-antimycotic with 2.5–10 µg/ml trypsin. When approximately 80% CPE was visible, cells and culture media were harvested, freeze/thawed twice and cell debris pelleted. Viral stocks were titrated by tissue culture infectious dose 50 (TCID_50_).

### 2.2. Reverse transcription and quantitative polymerase chain reaction

LLC-PK1 cells seeded into 6 well plates were mock infected or infected with PDCoV (103.8 TCID_50_ units/well). At the indicated time points, cells were scraped into PBS and were pelleted. Cell pellets were lysed in RLT buffer (Qiagen) and RNA extracted using an RNAeasy kit and following the manufacturer’s instructions. RNA was eluted into 50 µl RNAse-free water. Complementary DNA was generated using superscript IV (Invitrogen) following the manufacturer’s instructions and using 300 ng RNA and a random primer. Quantitative PCR was performed using Taqman Fast Universal 2x Master Mix (Invitrogen) including 125 nM probe, 500 nM primers and 2 µl cDNA in a 10 µl reaction. Primer and probe sequences within the PDCoV M gene have been described previously [34, 38] and absolute quantitation of cDNA copies was performed using a standard curve generated using a PCR product from the M gene covering the qPCR amplified fragment [34].

### 2.3. Western blot

LLC-PK1 cells seeded into 6 well plates were mock infected or infected with PDCoV (10^3.8^ TCID_50_ units/well). At the indicated time points, cells were scraped into PBS and pelleted. The cell pellet was lysed in 1x sample buffer (Biorad) containing β-mercaptoethanol, sonicated for 2 minutes (70% amplitude) and heated to 95 °C for 3 minutes. Proteins were separated by SDS-PAGE and transferred onto nitrocellulose membrane. After blocking in 5% milk in PBS-Tween 20, membranes were incubated with primary antibodies to detect PDCoV nucleoprotein (N) (Alpha Diagnostic International) and actin (Abcam) diluted in blocking buffer. After washing in PBS-T, membranes were incubated with IRDye labelled secondary antibody (LI-COR) diluted in blocking buffer. After further washes, membranes were imaged using an Odyssey CLx Infrared imaging system (LI-COR).

### 2.4. Virus growth curve and titration by TCID_50_

LLC-PK1 cells seeded into 6 well plates were mock infected or infected with PDCoV (10^3.8^ TCID_50_ units/well). At the indicated time points, culture media was harvested and stored at - 80 °C. Virus was titrated by TCID_50_, Briefly, cells seeded into 96 well plates were infected with a 2 fold serial dilution series of virus. Cells positive and negative for CPE were scored at 5 days post infection and viral titer calculated using the Reed and Muench method.

### 2.5. Immunofluorescence

LLC-PK1 cells seeded into 24 well plates on coverslips were mock infected or infected with PDCoV (10^3.3^ TCID_50_ units/well). At the indicated time points cells were fixed with 4% paraformaldehyde in PBS and permeabilized with 0.1% triton X-100 in PBS. After blocking in 0.5% bovine serum albumin (BSA) in PBS, cells were labelled with primary antibodies specific for PDCoV N and dsRNA (J2, English and Scientific Consulting Kft.). Cells were washed in PBS and labelled with Alexa Fluor conjugated secondary antibodies (Invitrogen) diluted in blocking buffer. After further washing, nuclei were stained using 4′,6-diamidino-2-phenylindole (DAPI), mounted onto glass slides using Vectashield (Vector Labs) and sealed using nail varnish.

For detection of nascent viral RNA, 30 minutes prior to fixation, cells were treated with 2 mM BrU (Sigma) and either 15 µM ActD (Sigma) or DMSO as a vehicle control. Cells were fixed and labelled as above with the following modifications; 0.1% fish skin gelatin in PBS was used as the blocking buffer and all steps following fixation were performed in an RNAse free environment with the inclusion of RNAsin (Promega) in all buffers to prevent loss of the BrU signal [39]. Primary antibody specific for BrdU, which also recognizes BrU, was purchased from Roche.

### 2.6. Transmission electron microscopy

LLC-PK1 cells seeded into 24 well plates with Thermanox coverslips (Thermo Fisher Scientific) were mock infected or infected with PDCoV (10^3.3^ TCID_50_ units/well). At the indicated time points, cells were fixed in 2% glutaraldehyde for 1 hour. Cells were incubated for 1 hour in 1% aqueous osmium tetroxide solution then dehydrated in increasing concentrations of ethanol. Following embedding in Agar 100 resin (Agar Scientific Ltd) and polymerisation overnight, 80 nm thick sections were cut, collected on hexagonal 200 thin bar copper grids and stained with lead citrate and 2% uranyl acetate. Data were collected using an FEI Tecnai 12 TEM at 100kV with a TVIPS F214 digital camera.

## 3. Results

### 3.1 Kinetics of porcine deltacoronavirus replication in LLC-PK1 cells

Initially, the dynamics of PDCoV OH-FD22 replication in LLC-PK1 cells were determined to both confirm successful completion of the virus replication cycle in these cells in our hands and to provide a context for subsequent experiments characterizing viral ROs. Accumulation of viral RNA was first measured by RT-qPCR using a primer-probe set specific for the M gene. This will, therefore, detect viral genomic RNA as well as all subgenomic RNAs containing the M gene sequence. Cells were infected or mock infected and at the indicated time points, RNA was extracted and RT-qPCR performed (Figure 1A). Viral RNA could be detected from the earliest time point tested (2 hours post infection (hpi)) but remained constant until 4 hpi. By 6 hpi, the total level of RNA had increased with a further increase detected by 8 hpi. This indicates synthesis of new viral RNA at these time points.

**Figure 1.**
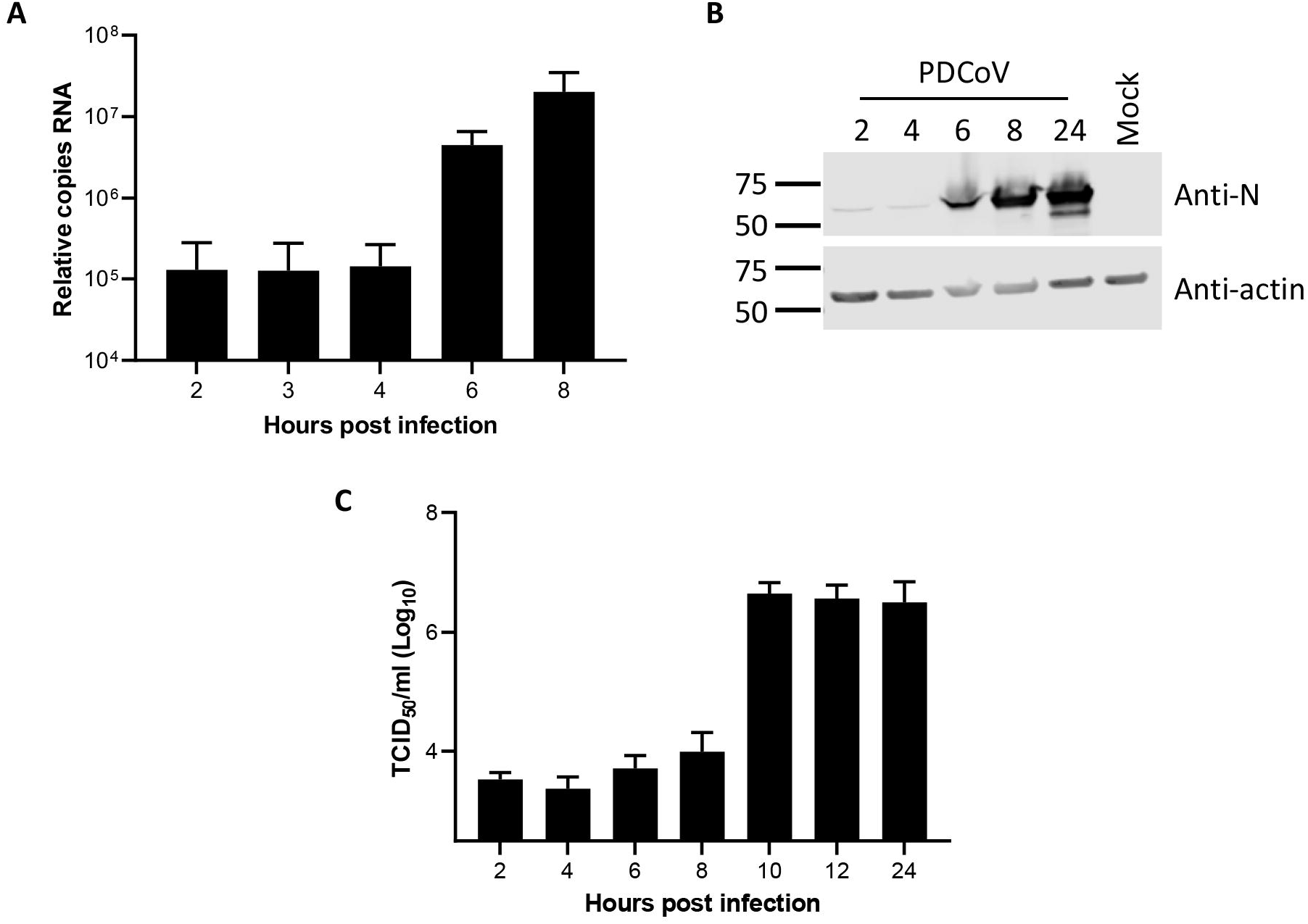
Dynamics of PDCoV OH-FD22 replication in LLC-PK1 cells. (**A**) LLC-PK1 cells were infected with PDCoV and at the indicated time points, RNA was harvested and reverse transcribed to cDNA. Copy number of cDNA was quantified by qPCR using a standard curve and normalized to mock infected cells. Mean and standard deviation of three independent replicates are shown. (**B**) LLC-PK1 cells were PDCoV infected or mock infected. Total cell lysate was harvested at the stated time points and viral nucleoprotein detected (Anti-N) by western blot. Actin (Anti-actin) was used as a loading control. Molecular weight markers are shown. Blot representative of three independent repeats. (**C**) LLC-PK1 cells were infected with PDCoV and at the indicated time points, cell culture media was harvested. The titer of progeny virus was determined by TCID_50_. Mean and standard deviation from three independent replicates are shown.

Following characterization of total viral RNA levels within infected cells, the levels of viral protein were measured. Cells were infected or mock infected and total cell lysates were harvested at the indicated time points. Presence of PDCoV N protein was then determined by western blot (Figure 1B). A band corresponding in size to PDCoV N was detected in all virus infected samples but not in the mock sample. A very low level of N could be detected at 2 and 4 hpi with an increasing level from 6-24 hpi. This confirms synthesis of new viral proteins from 6 hpi onwards.

Finally, release of progeny virions from PDCoV infected cells was measured. Cells were infected with PDCoV and cell culture medium harvested at the indicated time points. The amount of virus contained in these samples was measured by TCID_50_ (Figure 1C). Virus could be detected in all samples with a clear increase in the amount of virus present in the cell culture media from 10 hpi, indicating new virions are released from the host cell by this time point post infection. Taken together, this data confirms that PDCoV completes the full virus replication cycle in LLC-PK1 cells with synthesis of new viral RNA and protein occurring between 4 and 6 hpi and release of progeny virions between 8 and 10 hpi.

### 3.2 Visualizing porcine deltacoronavirus RNA synthesis

To begin to characterise PDCoV RO, the cellular location of N protein and dsRNA was visualized. Coronavirus replication is associated with the accumulation of dsRNA in cytoplasmic puncta [20, 24, 39, 40]. Although the precise nature and role of this dsRNA is debated, the location of dsRNA is often used as a marker for the sites of viral RNA synthesis. Therefore, cells were infected or mock infected and were fixed and labelled with antibodies specific for N and dsRNA at different time points following infection (Figure 2). No signal could be detected in mock infected cells or in cells fixed at 2 hpi. However, at 4 hpi, clusters of cytoplasmic puncta of N protein could be seen with a smaller number of dsRNA puncta also visible. These appeared in the same area of the cell, although did not appear to co-localize. By 6 hpi the number of both N and dsRNA puncta had increased. By 8 hpi, N signal was now found throughout the cytoplasm and appeared in a reticular staining pattern. The number of dsRNA puncta also continued to increase from 8 to 24 hpi. These results indicate that viral RNA synthesis is likely to begin from around 4 hpi.

**Figure 2.**
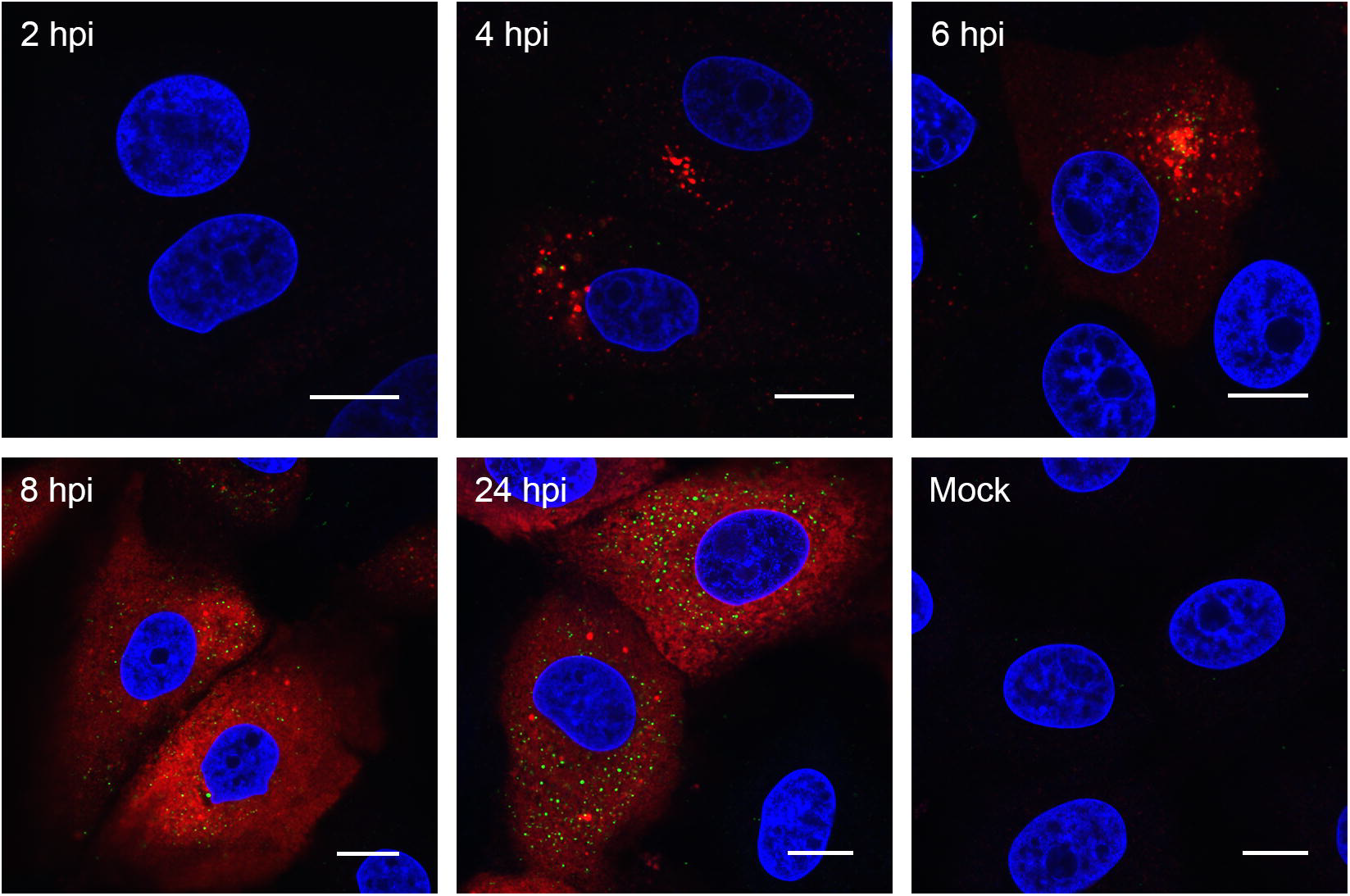
PDCoV associated dsRNA can be detected from 4 hpi. LLC-PK1 cells were infected or mock infected. At the stated time points, cells were fixed and labelled with anti-dsRNA (green) and anti-N (red). Nuclei were stained with DAPI (blue) and scale bar indicates 10 µm. Images are representative of three independent replicates.

To more conclusively determine the onset of viral RNA synthesis, as well as visualize where within the cell RNA synthesis is taking place, nascent RNA was labelled with 5-bromouridine (BrU). Cells were infected or mock infected and for 30 minutes prior to fixation, cells were treated with BrU in the presence of actinomycin D (ActD) to inhibit cellular RNA synthesis. BrU incorporated into nascent RNA was then detected using an anti-BrdU antibody (Figure 3). In mock infected cells incubated in BrU without ActD, cellular RNA was detected in both the nucleus and cytoplasm, as expected. In the presence of ActD, this signal was lost. This confirms that BrU signal detected in PDCoV infected cells is newly synthesized viral RNA. No BrU signal could be detected at 2 hpi. However, at 3 hpi, individual cytoplasmic puncta or small clusters of puncta could be seen. The number of puncta increased as infection progressed and they became more widely dispersed throughout the cytoplasm. This demonstrates that PDCoV RNA synthesis begins from 2.5-3 hpi in small clusters within the cytoplasm and by later time points, numerous sites of RNA synthesis exist within the cell.

**Figure 3.**
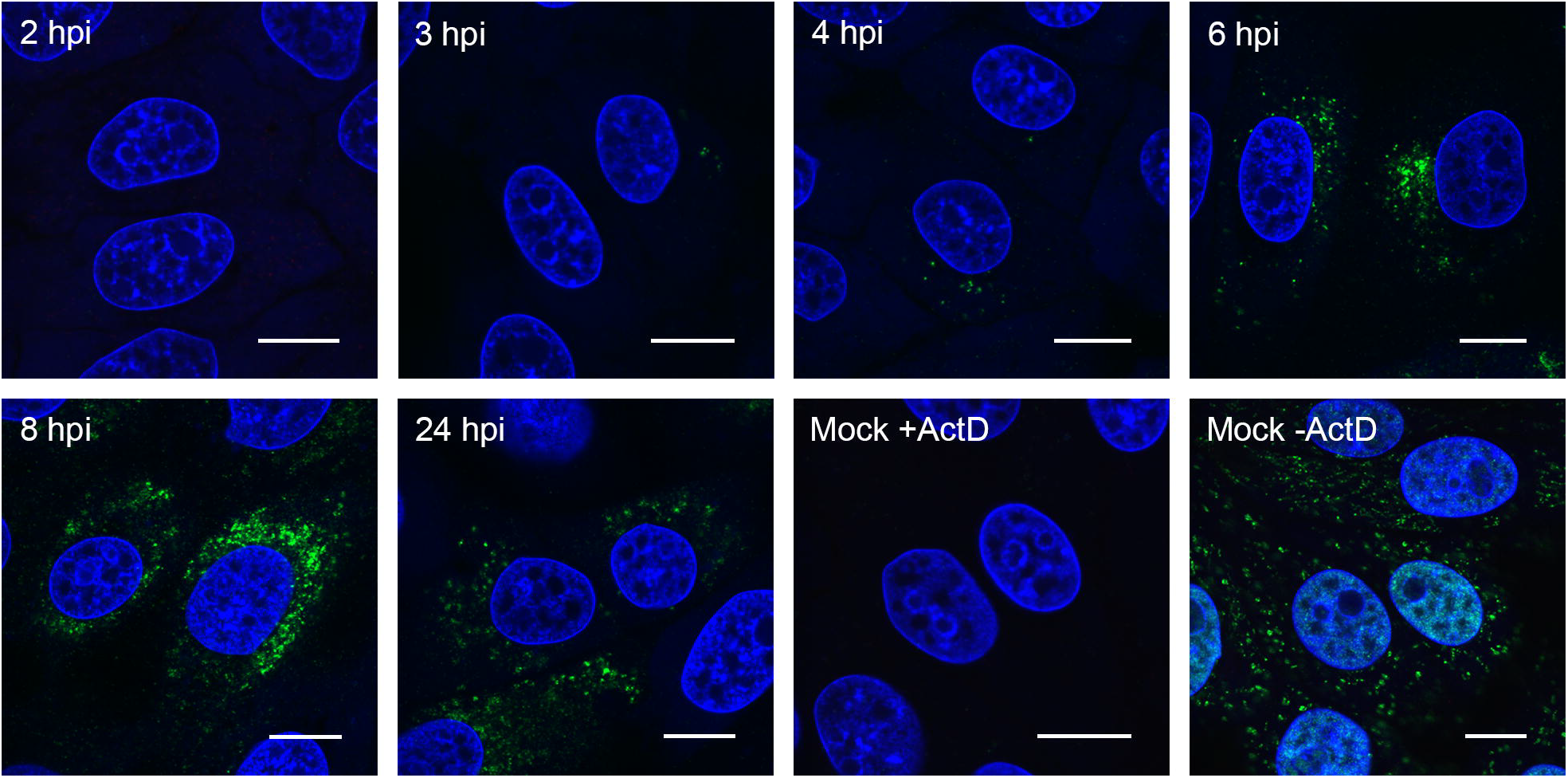
PDCoV RNA synthesis can be detected from 3 hpi. LLC-PK1 cells were infected or mock infected. Thirty minutes prior to the indicated fixation time, cells were incubated with 2 mM 5-Bromouridine (BrU) and 15 µM actinomycin D (ActD). Mock cells were incubated with (+ActD) and without (-ActD) actinomycin D as indicated. Cells were fixed and RNA containing BrU detected using an anti-BrdU antibody (green). Nuclei were stained with DAPI (blue), scale bar indicates 10 µm. Images representative of three independent replicates.

### 3.3 PDCoV ROs include double membrane vesicles and zippered ER with double membrane spherules

It has recently been reported that PDCoV infection triggers the formation of DMVs [36]. However, the presence of either CM or zippered ER and double membrane spherules has not been demonstrated. Therefore, a detailed analysis of PDCoV ROs was performed. Cells infected with PDCoV or mock infected were fixed at a range of time points post infection and were embedded and processed for transmission electron microscopy analysis. Initially samples from 8 hpi were imaged (Figure 4). At this time point virus particles in vesicles could be found, either as individual particles per vesicle or multiple particles in a single larger vesicle. In addition, DMVs were clearly visible (Figure 4). The samples were prepared by glutaraldehyde fixation and, as has been observed previously, under these conditions, DMVs had a fibrous content and the two membranes were closely apposed in some areas but in others had become separated from one another [21, 41, 42]. Large regions with numerous DMVs were observed as well as both individual DMVs and clusters of small number of DMVs. In addition to DMVs, regions of small double membrane vesicles interspersed with sections of paired membranes could be found. The small double membrane vesicles had very tightly apposed membranes and the membrane appeared to be lined with electron dense content. The areas of paired membranes and small double membrane vesicles were surrounded by electron density and appeared highly comparable to regions of zippered ER with associated spherules identified previously in cells infected with *Gammacoronavirus* IBV [24]. Both large and small regions of zippered ER and spherules were observed in PDCoV infected cells and they were found, most commonly, in the perinuclear region. Regions of zippered ER and spherules often, although not always, had a small number of DMVs in the vicinity. Together, this demonstrates that the PDCoV RO is made up of both DMVs and zippered ER with double membrane spherules.

**Figure 4.**
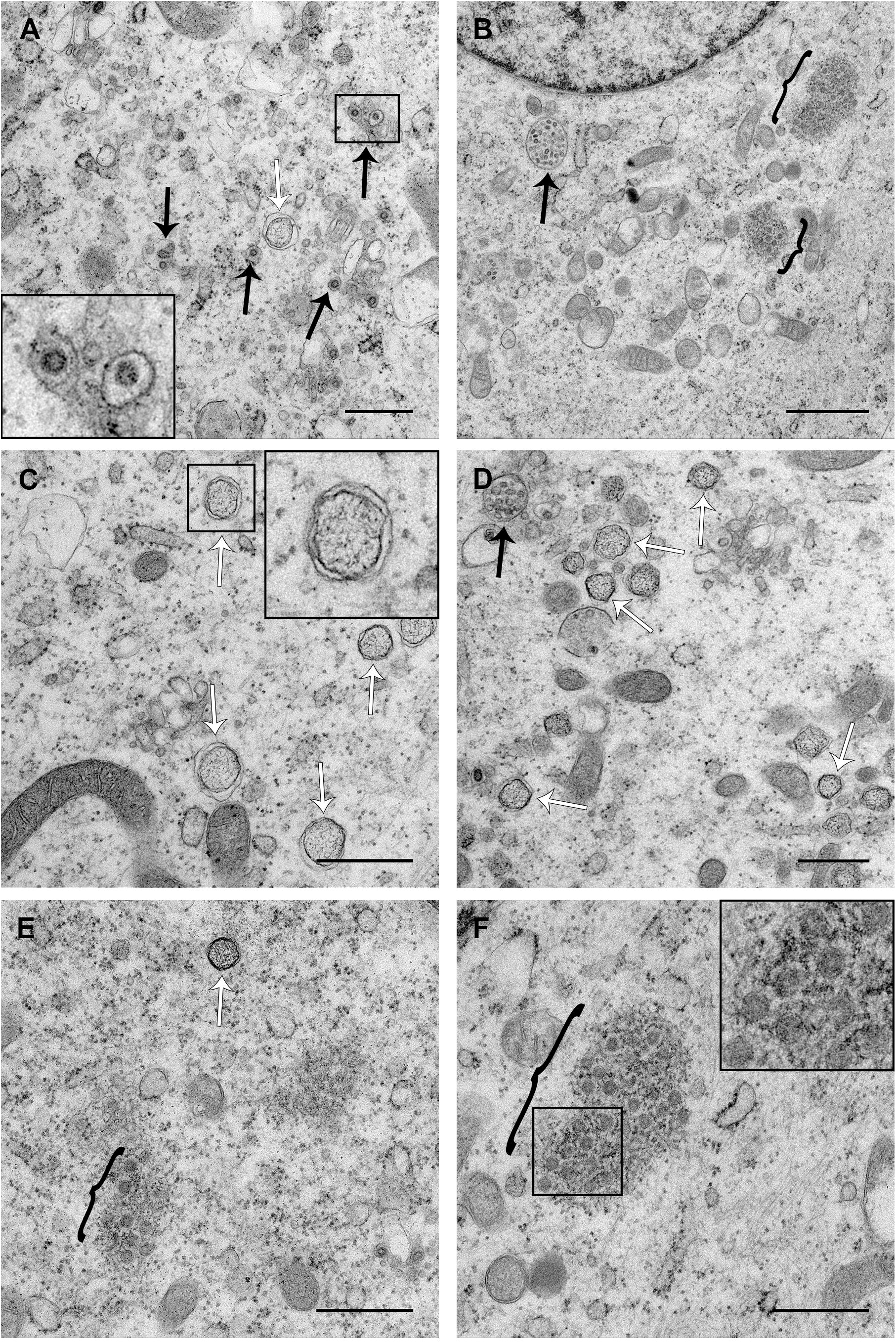
PDCoV RO is made up of DMVs and zippered ER with double membrane spherules. LLC-PK1 cells were infected with PDCoV. At 8 hpi, cells were fixed with glutaraldehyde and processed for transmission electron microscopy. Virions in vesicles are indicated with white arrows, DMVs are indicated with black arrows and regions of zippered ER with spherules are indicated with black brackets. Scale bars indicate 500 nm (**A** and **C-F**) or 1 µm (**B**).

### 3.4 PDCoV ROs, including zippered ER and double membrane spherules, are visible from 6 hpi

PDCoV RNA synthesis was detected in single puncta or small clusters of puncta from 3 hpi, larger clusters of puncta from 4 hpi and by 6hpi, the number of puncta had dramatically increased (Figure 3). Therefore to investigate further whether ROs associated with this RNA synthesis could be detected earlier in infection than 8 hpi, a range of time points from 4–24 hpi were imaged (Figure 5). No ROs or any other evidence of virus infection were detected at 4 hpi. However, at 6 and 24 hpi, ROs comprising both DMVs and regions of zippered ER and double membrane spherules were found. ROs visualized at both 6 and 24 hpi were highly comparable to those seen at 8 hpi. In addition, at 24 hpi, as well as virus particles in vesicles, virus particles were observed in the ER, presumably as a result of budding into the ER. Together this confirms the presence of ROs made up of DMVs, zippered ER and spherules from 6 to 24 hpi, during the peak of PDCoV RNA synthesis.

**Figure 5.**
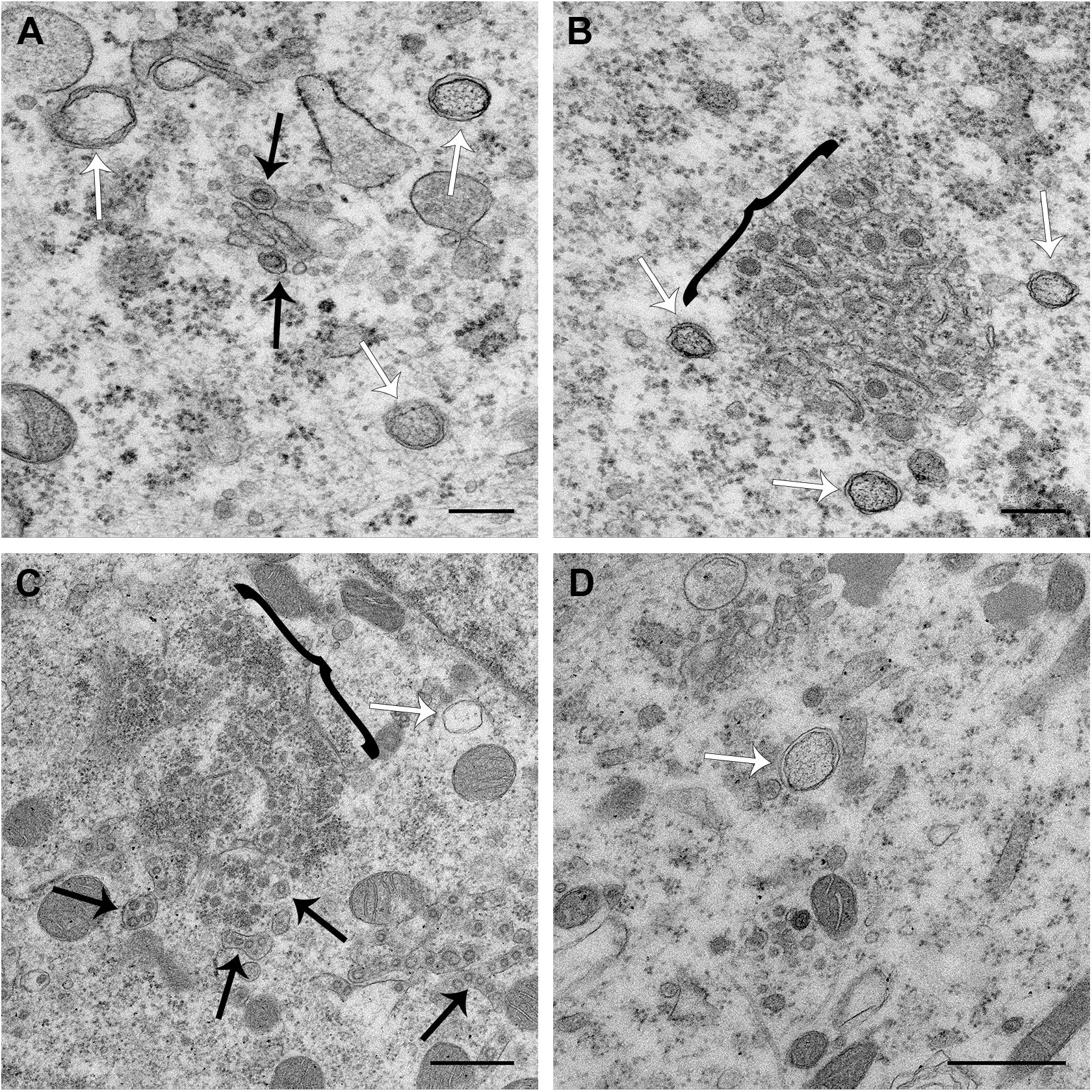
DMVs and zippered ER with double membrane spherules are present from 6 to 24 hours post infection. LLC-PK1 cells were infected with PDCoV and at 6 (A and B) or 24 (C and D) hours post infection, cells were fixed with glutaraldehyde and processed for electron microscopy. Virions in vesicles are indicated with white arrows, DMVs are indicated with black arrows and regions of zippered ER with spherules are indicated with black brackets. Scale bars indicate 200 nm (**A**) or 500 nm (**B-D**).

## 4. Discussion

In this study, we have characterised the replication dynamics of PDCoV OH-FD22 in porcine LLC-PK1 cells, with a particular focus on RO formation. Initial characterization of viral RNA, protein and progeny production demonstrated that synthesis of viral RNA, with an increase in accumulated RNA levels, could be detected from 6 hpi. This corresponded with detection of increased levels of viral protein at the same time point. Subsequent release of progeny virus was detected from 10 hpi. This data is broadly consistent with observations for the replication dynamics of other coronaviruses, including MHV, SARS-CoV, IBV and another lab-adapted strain of PDCoV [21, 24, 36, 43–45].

Following on from initial characterization of viral replication dynamics, the accumulation of virus replication associated dsRNA was visualized. Although historically used as a marker for sites of RNA synthesis, the role of dsRNA during coronavirus infection is not currently clear. It localizes to the interior of DMVs in SARS-CoV infected cells [20] and although it co-localizes with sites of active RNA synthesis at early time points in MHV infected cells, this is less obvious at later time points [39]. In addition, there is only partial co-localization between dsRNA and replicase proteins from other coronaviruses and more distantly related nidoviruses [5, 20, 21, 41, 42, 46, 47]. Therefore, whether dsRNA represents a *bone fide* intermediate in viral RNA synthesis, is a byproduct of virus replication or performs some other function is not currently understood. Despite this, dsRNA accumulation does correlate with the onset of viral RNA synthesis and provides a useful marker. Therefore, accumulation of dsRNA in PDCoV infected cells and co-localization with N protein was determined. Both dsRNA and N were detected in a small number of cytoplasmic puncta from 4 hpi and these puncta were clustered together. From 6 hpi, the number of dsRNA puncta increased and the puncta became more dispersed throughout the cytoplasm, as has been seen for other coronaviruses [20, 24, 39]. The number of N puncta also increased by 6 hpi and from 8 hpi onwards, N signal predominantly showed a cytoplasmic reticular pattern. This demonstrates that the onset of PDCoV protein synthesis occurs prior to 4 hpi, earlier than was detected by western blot. It is not surprising that small changes in N levels are less likely to be detected at the population level as determined by western blot than when imaging individual infected cells. Also in agreement with previous observations, there appeared to be little co-localization between dsRNA and N at any time point studied here.

To gain a more conclusive picture of the onset and localization of viral RNA synthesis, BrU incorporation into nascent viral RNA was adopted. This allows the visualization of RNA synthesized within a defined period of time, here 30 minutes prior to fixation of cells. Using this technique, the onset of viral RNA synthesis was shown to be between 2.5 and 3 hpi. Again, this is significantly earlier than was detected at the population level using RT-qPCR. Sites of viral RNA synthesis appeared in individual puncta or small clusters of puncta in the perinuclear region. As infection proceeded, comparable to sites of dsRNA accumulation, the number of BrU puncta increased and they became more dispersed throughout the cytoplasm. By 4 hpi there was a small increase in the number of puncta but the number and spread of puncta increased dramatically by 6 hpi. Therefore, it would be expected that there are a small number of ROs at 3 and 4 hpi but by 6 hpi, numerous ROs would be present throughout large areas of the cytoplasm. The dramatic increase in the number of sites of RNA synthesis (BrU puncta) by 6 hpi correlates with the observed increase in accumulated viral RNA detected by RT-qPCR, indicating that RT-qPCR is not sufficiently sensitive to detect small changes in total RNA at earlier time points.

Finally, the ultra-structure of PDCoV ROs was investigated. In agreement with previous studies on PDCoV and numerous other coronaviruses, DMVs were found in PDCoV infected cells. Furthermore, the appearance of these DMVs was indistinguishable from DMVs found in other coronavirus infected cells when fixed with glutaraldehyde. DMVs were first detected at 6 hpi and continued to be detected until 24 hpi, the latest time point studied. Throughout infection, DMVs were observed as individual vesicles or in either small or large clusters. However, as was seen for IBV [24], the DMVs were rarely in close proximity making it unlikely that they would have connections and form part of an interconnected membrane network, as was observed for SARS-CoV [20]. In addition, the large clusters of DMVs were not found to be associated with zippered ER and spherules suggesting there may be functional differences between these structures. Finally, no likely intermediates in DMV formation were observed at any time point. Therefore, it remains to be determined precisely how DMVs are formed.

The most significant finding from this study is the presence of, in addition to DMVs, regions of zippered ER and associated double membrane spherules in PDCoV infected cells. Furthermore, no regions of more branching CMs were identified. Zippered ER and spherules were detected from 6 to 24 hpi and their appearance did not alter over this time course. Throughout infection, both large and small areas of zippered ER and spherules were found. These were predominantly in the perinuclear region and often, although not always, a small number of DMVs were found in the proximity. Significantly, the regions of zippered ER and spherules were highly comparable to those identified previously by us in IBV infected cells [24]. Therefore, the ROs of *Gamma*- and *Deltacoronaviruses* appear to be conserved.

Despite the observed difference in the ROs induced by different coronavirus genera, the role of the different parts of the RO during coronavirus replication and the precise location of viral RNA synthesis on these membranes remain key questions in the field. Understanding the role of the different parts of the RO will shed light on the significance of the differences observed between the structures induced by the different coronavirus genera. Ultimately, understanding the function and formation of coronavirus ROs will provide useful information to allow control of this important virus family.

## Acknowledgments

Many thanks to Prof. Linda Saif, The Ohio State University, for kindly sharing porcine deltacoronavirus OH-FD22 and providing advice on protocols for virus propagation. Thank you also to Paul Britton and Pip Beard for helpful discussions. This research was funded by Biotechnology and Biological Sciences Research Council, grant numbers BB/N002350/1, BBS/E/I/00007034, BBS/E/I/00007037, BBS/E/I/00007038 and BBS/E/I/00007039.

